# Photoactivated SOPP3 enables APEX2-mediated proximity labeling with high spatio-temporal resolution in live cells

**DOI:** 10.1101/2024.06.23.595995

**Authors:** Dajun Qu, Yaxin Li, Qian Liu, Biao Cao, Mengye Cao, Peng Zou, Hu Zhou, Wenjuan Zhang, Weijun Pan

**Affiliations:** CAS Key Laboratory of Nutrition, Metabolism and Food Safety, State Key Laboratory of Experimental Hematology, Shanghai Institute of Nutrition and Health (SINH), University of Chinese Academy of Sciences, Chinese Academy of Sciences, Shanghai, 200031, China; Department of Analytical Chemistry, State Key Laboratory of Drug Research, Shanghai Institute of Materia Medica, Chinese Academy of Sciences, Shanghai, 201203, China; College of Chemistry and Molecular Engineering, Synthetic and Functional Biomolecules Center, Beijing National Laboratory for Molecular Sciences, Key Laboratory of Bioorganic Chemistry and Molecular Engineering of Ministry of Education, Peking University, Beijing, 100871, China

**Author notes:** These authors contributed equally: Dajun Qu, Yaxin Li, Qian Liu. Corresponding to: Wenjun Pan, Wenjuan Zhang and Hu Zhou.

**Keywords:** proximity labeling, photosensitizer, APEX2, SOPP3, protein interactome

## Abstract

Protein interactome characterized via proximity labeling (PL) and proteomic analysis is important for mechanism study in biomedical research. However, exogenous H_2_O_2_ required for engineered peroxidase, like APEX2, is cytotoxic, while biotin ligases-based PL have poor temporal resolution. Here, we showed that SOPP3, an engineered photosensitizer, could trigger APEX2 mediated PL within seconds under blue light illumination, without exogenous H_2_O_2_ or cytotoxicity. We demonstrated that APEX2 plus photoactivated SOPP3 could catalyze spatially restricted biotinylation at endoplasmic reticulum-mitochondria contact sites and cell-cell interface. Through fusing APEX2 and SOPP3 as a chimeric probe, we showed that APEX2-SOPP3 also works well in multiple subcellular locations, including cell surface. Notably, either APEX2 plus SOPP3, or chimeric APEX2-SOPP3 requires 10 folds lower illumination power, compared to that for light-oxygen-voltage protein alone mediated PL, and has better response to illumination in terms of PL in time course with little cytotoxicity. Thus, this new PL technology paves an avenue to characterize molecular dynamics of key biomedical events with high spatial-temporal resolution, efficiency, and flexible applicability.

## Main

Proximity labeling (PL) is an emerging technology for probing protein interactome in living cells. Current PL technology, like engineered peroxidases (APEX2, HRP), are widely employed, benefiting from their high temporal resolution^1,2^. However, cytotoxicity of exogenous H_2_O_2_, used to activate APEX2, largely limits its application. Meanwhile, biotin ligases, such as BioID^3^, BioID2^4^, TurboID^5^, give poor temporal resolution due to low catalytic activity and prolonged labeling time. An approach that could catalyze PL in living cells with high spatio-temporal resolution is still missing.

To address this gap, we employed light, which can be flexibly controlled in most biomedical experiments with high temporal and spatial precision, as a trigger for PL reaction. Singlet oxygen photosensitizing protein-3 (SOPP3) (∼12 kDa), a member of light-oxygen-voltage (LOV) family, has been validated as a genetically encoded photosensitizer^6^, which produces a high level of H_2_O_2_ in the presence of superoxide dismutase (SOD) under blue light illumination^7,8^. Given the ubiquitous presence of SOD in cells^8^, we hypothesized that SOPP3 under the control of blue light illumination might enable APEX2 to oxidize biotin-phenol (BP) into reactive phenoxyl radicals, which preferentially labels proximal proteins in living cells without exogenous H_2_O_2_ (Fig. 1a).

**Figure 1.**
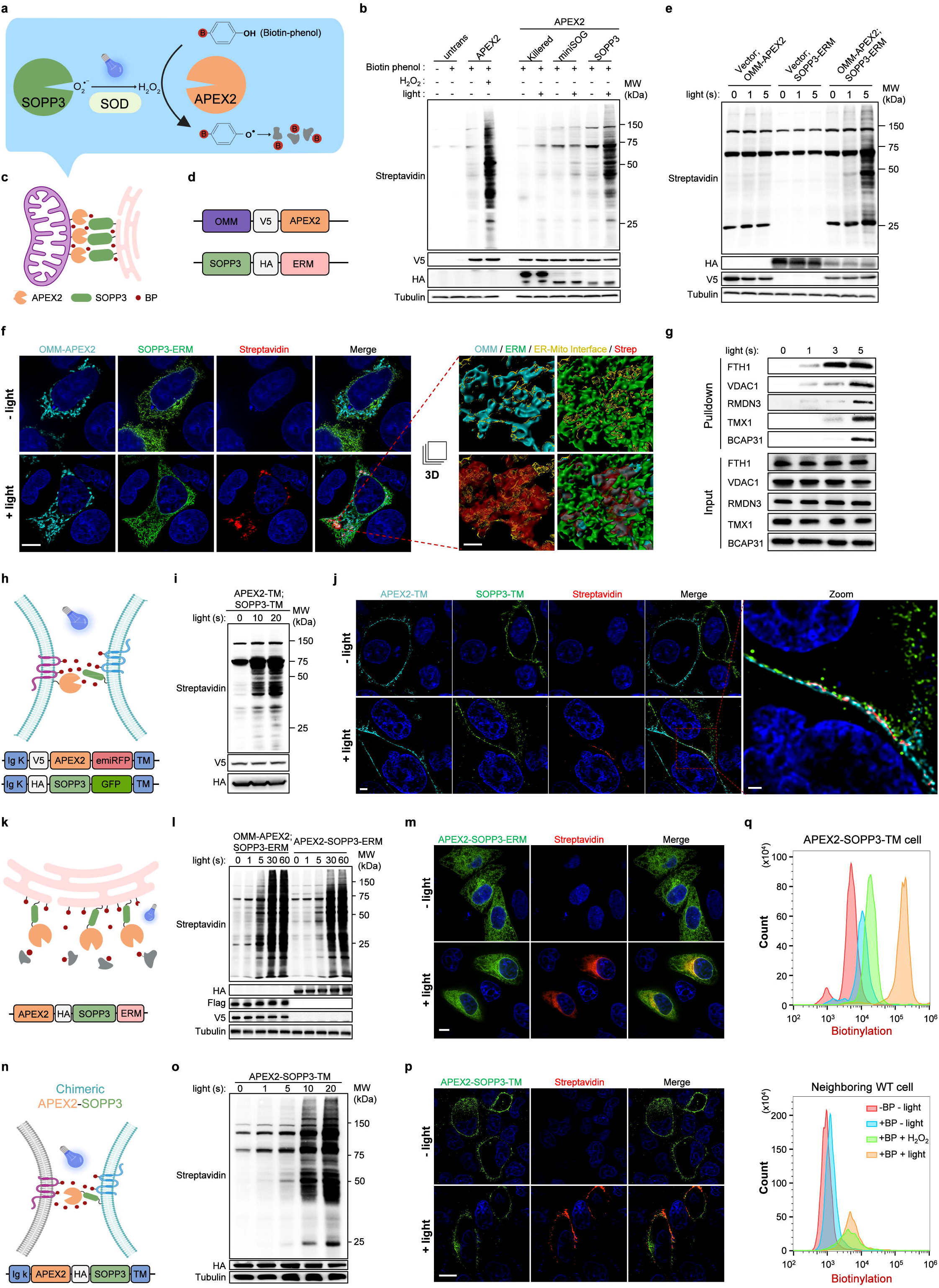
Photoactivated SOPP3 triggers APEX2-mediated proximal labeling with high spatio-temporal resolution. **a** Schematic of APEX2+SOPP3 mediated photoactivated-proximal labeling (photo-PL). B, Biotin. **b** Photosensitizer screening for APEX2 activation. Streptavidin blot showed photo-PL efficiency. anti-V5 and anti-HA indicated expression level of APEX2 and photosensitizers, respectively. Anti-α-tubulin, internal loading control for western blot. **c**,**d** Schematic (**c**) and construct designs (**d**) of APEX2+SOPP3 mediated PL to map proteome on ER-Mito contact sites. BP, Biotin Phenol. **e** Evaluation on the efficiency of photo-PL at ER-Mito contact sites. Anti-HA and anti-V5 indicated expression level of SOPP3-ERM and OMM-APEX2 respectively. **f** Left: confocal fluorescence imaging of APEX2+SOPP3 mediated photo-PL on ER-Mito contact sites. Outer membrane of mitochondria (OMM)-localized APEX2 and ER membrane (ERM)-localized SOPP3 were visualized by anti-V5 and anti-HA antibody. Biotinylation signals on the contact sites were visualized by Alexa Fluor 555-conjugated streptavidin. Illumination time, 5 seconds. Scale bar, 9 μm. Right: the ‘Surface’ tool in Imaris software was used to create a 3D rendering from each channel of confocal images of boxed region. Contact area algorithm was further performed to determine the interface (yellow) between ER (green) and Mito (cyan). Scale bar, 0.8 μm. **g** Validation on MAMs proteins from streptavidin-enriched PL samples. **h**,**i** Schematic and construct designs (**h**) and evaluation on photo-PL efficiency on cell-cell contact sites via western blotting analysis (**i**). Anti-V5 and anti-HA indicated expression level of APEX2-TM and SOPP3-TM respectively. **j** Confocal fluorescence imaging of APEX2+SOPP3 mediated photo-PL on cell-cell contact sites, stained with Alexa Fluore 555-conjugated streptavidin. Scale bar, 2 μm. Zoomed image from boxed region showed biotinylation labeling precisely on cell-cell contact sites. Scale bar, 1 μm. Illumination time, 10 seconds. **k** Schematic and construct design of chimeric APEX2-SOPP3-mediated photo-PL in ERM. **l** Evaluation on the efficiency of photo-PL mediated by chimeric APEX2-SOPP3 targeted to ERM via western blotting analysis. Anti-HA/Flag/V5 indicated expression level of chimeric APEX2-SOPP3-ERM, SOPP3-ERM and OMM-APEX2 respectively. **m** Confocal fluorescence imaging of photo-PL via APEX2-SOPP3 targeted to ERM. Chimeric APEX2-SOPP3-ERM and biotinylation signals was visualized by anti-HA antibody and Alexa Fluor 555-conjugated streptavidin respectively. Anti-Flag blot indicated expression level of chimeric APEX-SOPP3 targeted to various subcellular compartments. Illumination time, 5 seconds. Scale bar, 10 μm. **n**,**o** Schematic and construct designs (**n**) and evaluation on chimeric APEX2-SOPP3 on cell-cell contact sites via western blotting analysis (**o**). Anti-HA indicated expression level of chimeric APEX2-SOPP3-TM. **p** Confocal fluorescence imaging of chimeric APEX2-SOPP3 mediated photo-PL on cell surface. Chimeric APEX2-SOPP3 was visualized by anti-HA antibody. Biotinylation signals were visualized by Alexa Fluor 555-conjugated streptavidin. Scale bar, 8 μm. Illumination time, 5 seconds. **q** Flow cytometry analysis on **p** (30,000 cells per condition). After photo-PL, cells were stained with Alexa Fluor 555-conjugated streptavidin, followed with flow cytometry gating for quantification of biotinylation labeling on the surface of APEX2-SOPP3-TM cells (anti-HA positive, **q** up) and neighboring WT cells (anti-HA negative, **q** down). Blue light-activated SOPP3 showed more efficient to facilitate APEX2-mediated proximity labeling compared to that of 1 mM H_2_O_2_ treatment. Illumination time, 10 seconds.

To test this hypothesis, we generated SOPP3 expression construct, alone with other photosensitizers (KillerRed and miniSOG), and transfected them with APEX2 into HeLa cells. SOPP3, plus APEX2, indeed exhibited unique PL activity, while APEX2 or SOPP3 alone showed little PL activity under the same illumination condition (Fig. 1b, Supplementary information, Fig. S1a). Moreover, we showed that SOD was indispensable for APEX2+SOPP3 mediated PL labeling (Supplementary information, Fig. S1b-c). To verify SOPP3-assisted biotinylation activity of APEX2 is proximity-dependent, we employed FRB-FKBP interaction system^9^ to control the proximity between APEX2 and SOPP3 via rapamycin induction. The Mitochondrial outer membrane (OMM) localized APEX2 (facing cytosol, signal peptide of OMM protein-TOM20) and cytosol resident SOPP3 were conjugated with FKBP and FRB respectively. Streptavidin blot showed that biotinylation activity via SOPP3+APEX2 was dramatically increased with rapamycin treatment (Supplementary information, Fig. S1d-e). Thus, we concluded that photoactivated SOPP3 could trigger APEX2-mediated PL. Importantly, a marked amount of biotinylation could be achieved in 5 seconds with biotinylation peaking at 20 seconds (Supplementary information, Fig. S1f). More important, the PL process showed little cytotoxicity (Supplementary information, Fig. S1g-h) and terminated when the blue light illumination stopped (Supplementary information, Fig. S1i).

To demonstrate the applicability of this new PL technology, we tested OMM-APEX2 and SOPP3-ERM (fused with ER membrane resident protein-SEC61B, facing cytosol) mediated PL on Endoplasmic reticulum (ER)-mitochondria (Mito) contacts sites (Fig. 1c-d). Streptavidin blot also showed strong biotinylation signals within 5 seconds illumination, while either OMM-APEX2 or SOPP3-ERM alone showed little biotinylation activity under the same illumination condition (Fig. 1e). Confocal imaging analysis showed high spatially restriction of biotinylation signal at the ER-Mito interface with 5 seconds blue light illumination (Fig. 1f, Supplementary information, Video S1).

To test whether APEX2+SOPP3-mediated PL could uncover proteins enriched on ER-Mito contact sites, we carried out tandem mass tag (TMT) labeling-based quantitative proteomics experiment on our streptavidin enriched sample and applied two sequential filtering to analyze proteomic data (Supplementary information, Fig. S2-S3 and Table S1-2). Proteome list of 84 proteins was finally generated (Supplementary information, Fig. S4a), which showed strong ER and mitochondrial membrane association after Gene Ontology Cellular Component (GOCC) analysis (Supplementary information, Fig. S4b), indicating high spatial resolution of this APEX2+SOPP3 mediated PL system. Notably, listed proteins included 13 well-characterized mitochondria-associated membrane (MAM) proteins (e.g., VDAC1, TMX1, SAR1A, RMDN3, HSPA9, GDAP1, DNM1L, CLCC1, CISD1, CANX, AKAP1, BCAP31 and C1QBP), and have more proteins related to typical MAM functions, such as FUNDC2 (autophagy) and FIS1 (mitochondrial fission regulation) (Supplementary information, Fig. S4a). And this dataset is comparable to previous MAM proteomes revealed by Split-TurboID^10^ or Contact-ID^11^ (Supplementary information, Fig. S4c).

We then carried out western blotting validation on streptavidin-enriched biotinylated protein after APEX2+SOPP3 mediated PL in time course. VDAC1, RMDN3, TMX1 and BCAP31, as identified in our and previously reported proteome, could be found in streptavidin-enriched PL samples with 5 seconds blue light illumination (Fig. 1g). Surprisingly, VDAC1, RMDN3 (two well-known OMM proteins) and a novel ER-Mito contact-associated protein, ferritin heavy chain1 (FTH1), could be detected with only 1 second illumination (Fig. 1g). Meanwhile, TMX1 and BCAP31 (two well-known ERM proteins) could be detected with only 3-5 seconds illumination (Fig. 1g). The association of FTH1 with mitochondria was further confirmed via Proteinase K protection assay on the MAM of crude mitochondria (Supplementary information, Fig. S4d).

Cell-cell interactions widely existed in multicellular organisms, from transient interaction among motile immune cells, to long standing intercellular contacts in epithelia^12^. Molecular characterization on cell-cell interface is important to understand the underlying mechanism of cell-cell adhesion, recognition, signaling transduction etc. Encouraged by APEX2+SOPP3 mediated PL efficiency in ER-Mito contact sites, we next explored whether APEX2+SOPP3 mediated PL could be achieved on cell-cell interface. We established stable cell lines expressing cell surface resident APEX2 or SOPP3 respectively, and then mixed them equally for PL reaction (Fig. 1h). After BP incubation and subsequently blue light illumination, protein biotinylation were confirmed by western blot (Fig. 1i). Further, confocal imaging analysis showed that biotinylation signal was precisely enriched on the interface of neighboring cells, with surface expression of APEX2 or SOPP3 respectively (Fig. 1j, Supplementary information, Video S2). Compared with other split-version of PL methods, including split-APEX2^13^, split-TurboID^10^ or Contac-ID^11^, we found that photoactivated APEX2+SOPP3 system dramatically improves temporal resolution (less than 5 seconds) of PL in living cells with high efficiency and reversible control (Supplementary information, Fig. S4e).

To increase the versatility of this method, we fused APEX2 and SOPP3 as a chimeric protein (Fig. 1k). As expected, chimeric APEX2-SOPP3 targeted to ERM also showed strong PL activity and clear spatial restriction (Fig. 1l-m; Supplementary information, Fig. S5a). In addition, chimeric APEX2-SOPP3 expressing in nucleus, ER lumen, outer mitochondrial membrane, or mitochondrial matrix all worked well (Supplementary information, Fig. S5c-d). Thus, we explored whether the chimeric APEX2-SOPP3 could label cell-cell interface. After establishment of stable cell line with chimeric APEX2-SOPP3 expression on cell surface (Fig. 1n), we mixed them with parent/wildtype (WT) cells for PL reaction under blue light or exogenous H_2_O_2_ induction. Biotinylation activity were detected by western blot and confocal imaging further showed strong biotinylation on the surface of APEX2-SOPP3-TM expressing cells under blue light illumination (Fig. 1o-p; Supplementary information, Fig. S5b). Flow cytometry analysis further confirmed that APEX2-SOPP3-TM, like exogenous H_2_O_2_, mediated biotinylation on the surface of parent/WT neighboring cells. Surprisingly, our data showed that 10-seconds illumination was more efficient than that of 1 mM exogenous H_2_O_2_ in terms of PL efficiency on the surface of APEX2-SOPP3-TM expressing cells (Fig. 1q). Taken together, our data indicated that chimeric APEX2-SOPP3 is also a useful tool to perform PL exploration via tagging to various subcellular localization, including cell-cell interface.

LITag is recently reported as a blue light illumination-based PL approach^14^. However, we found high power illumination (450 mW/cm^2^) required for LITag activation could directly cause severe mitochondria damage and cell death (Supplementary information, Fig. S6a-b), and LITag-mediated PL showed much lower biotinylation activity within 10 seconds illumination (Supplementary information, Fig. S6c). In contrast, SOPP3-enabled APEX2-mediated PL only requires mild illumination (50 mW/cm^2^), which is safe and responses well to illumination in time course (Supplementary information, Fig. S6a-c). Improved biotin ligase-based PL system, like PhastID, achieved protein interactome characterization in 15 min, while LOV-Turbo enable PL reaction under the control of blue light illumination, both has no H_2_O_2_ requirement^15,16^. However, compared with those methods, 5 seconds labeling window of APEX-SOPP3 with flexible control enable capturing of more precise molecular dynamics in living cells (Supplementary information, Fig. S6d).

In summary, SOPP3-enabled APEX2 activation under the control of mild blue light illumination is a novel and advanced PL method without exogenous H_2_O_2_ or cytotoxicity. SOPP3-APEX2 mediated PL reaches high spatio-temporal resolution, super efficiency, and flexible applicability, which would be a powerful method for mechanism studies via the exploration and characterization of interactomes in various biomedical research.

## Materials and Methods

### Plasmid Construction

All constructs were generated by oligos as listed in Supplementary Table 3, with standard cloning techniques. PCR fragments were amplified using KOD polymerase (TOYOBO). Vectors were digested using enzymatic restriction digestion and ligated to gel purified PCR products using Seamless Cloning Kit (Beyotime). Ligated plasmid products were transformed into competent TOP10 bacteria (TIANGEN) and confirmed by DNA sequencing.

### Cell culture, plasmid transfection and stable line generation

HeLa and HEK293T cells were maintained in Dulbecco’s modified Eagle’s medium (DMEM, Gibco) with 10% Fetal Bovine Serum (FBS) and Antibiotic-Antimycotic (1:100, Gibco). All cells were cultured in 37°C incubator with 5% CO_2_. Plasmid transfection was carried out with Lipofectamine 3000 (Invitrogen, L3000075) according to manufacturer’s instruction. To establish stable cell line used in this study, lentivirus containing pLVX-APEX2-TM, pLVX-SOPP3-TM, pLVX-APEX2-SOPP3-TM were produced respectively, using pLVX/psPAX2/pMD2G packaging system in HEK293T cells. HeLa cells were infected, followed with puromycin selection.

### Cell viability assay

Cell viability was measured with CCK-8 Assay Kit (MCE, HY-K0301) following the manufacturer’s instructions. 10,000 cells were seeded in 96-well plates for 24 h and then subjected to illumination for indicated time. After illumination, each well was added 10 μL of CCK-8 solution, and then incubated at 37°C for 2 h, followed with measurement of the absorbance at 450 nm by a microplate reader at indicated time point.

### APEX2+SOPP3 mediated PL

For APEX2+SOPP3 PL in HeLa cells, 500 μM Biotin phenol (BP) (TargetMol, T19209) solution was added and the cells were incubated at 37°C for 60 min. Labeling was triggered by a blue LED lamp (3S-Tech, https://www.3s-tech.net/products/ilmt.html) for indicated time at 30 cm directly above the cells. Light power was 50 mW/cm^2^ (measured by BENUO OPTICS HA350). Subsequently, labeling solution was quickly aspirated and cells were washed five times with ice cold quencher solution (10 mM sodium azide, 10 mM sodium ascorbate and 5 mM Trolox). Further assays were carried out after the extensively washes.

### Small molecule compound treatment

To induce FRB-FKBP proximity, cells were transfected with OMM-V5-FKBP-APEX2 and FRB-FLAG-SOPP3-NES. After transfection, cells were treated with rapamycin (TargetMol, T1537) for indicated time. For SOD1 inhibitor treatment assay, cells were transiently transfected with V5-APEX2-NES and HA-SOPP3-NES, then treated with ATN-224 (MCE, HY-16074) or LCS-1 (MCE, HY-115445) for 16 h. Until the last hour of treatment, 500 μM BP was added to culture medium before illumination to trigger APEX2+SOPP3 mediated PL.

### Streptavidin beads pulldown

Streptavidin pulldown was performed as previously descried^17^. Streptavidin agarose beads (Millipore, S1638) were washed three times with RIPA lysis buffer and incubated with clarified cell lysates at 4°C overnight. On the subsequent day, beads were then washed twice with RIPA lysis buffer, once with 1 M KCl, once with 0.1 M Na_2_CO_3_, once with 2 M urea in 10 mM Tris-HCl (pH=8.0), and twice with RIPA lysis buffer again. For western blotting analysis, proteins were denatured and eluted by incubation in SDS sample buffer with 2 mM biotin at 98°C for 5 min. For proteomic analysis, the beads were resuspended in washing buffer (75 mM NaCl in 50 mM Tris-HCl pH 8.0).

### On-bead trypsin digestion of biotinylated proteins

Proteomic samples for mass spectrometry analysis were prepared as previously reported^10,18^. In brief, 60 μL streptavidin beads-enriched samples were subjected to a double wash with 200 μL of 50 mM Tris-HCl (pH 7.5), which was then followed by two additional rinses with 200 μL of 2 M urea in 50 mM Tris-HCl (pH 7.5). After washing, samples were incubated with 80 μL of 2 M urea in 50 mM Tris-HCl (pH 7.5) containing 1 mM DTT and 0.5 μg trypsin (Promega) at 25°C with shaking for pre-digestion. After 1 hour, the supernatant was transferred to a fresh tube while the beads were shaken with the same buffer used for digestion for 30 minutes. The supernatant was combined with the previous elution. The remaining beads were double washed with 60 μL of 2 M urea in 50 mM Tris-HCl (pH 7.5), and the washing solutions were combined with the on-bead digest supernatant. The resulting mixture was thereafter reduced with 4 mM DTT for 30 minutes at 25°C with shaking, and subsequently alkylated with 10 mM iodoacetamide for 45 minutes in the dark at 25°C with shaking. An additional 1 μg of trypsin was added to each sample, which was then incubated at 25°C with shaking. After overnight digestion, the samples were acidified by adding formic acid (FA) to a final concentration of 1% FA (pH<3). Then, the samples were desalted on C18 StageTips and dried using Speed-Vac apparatus (Thermo Fisher Scientific).

### Tandem mass tag (TMT) labeling and fractionation of peptides

Desalted peptides were labeled with TMT 10-plex reagents (Thermo Fisher Scientific). Dried peptides for each sample were dissolved in 100 μL of 50 mM TEAB and each 0.8 mg vial of TMT reagent dissolved in 41 μL of anhydrous acetonitrile was added. After 1-hour incubation at room temperature, 8 μL of 5% hydroxylamine was added to quench the labeling reaction for 15 min at room temperature. Afterwards, all labeled peptides were pooled together, dried down via Speed-Vac, and subsequently desalted on a reversed phase tC18 SepPak column (Waters).

The TMT labeled peptides were fractionated using high pH reversed-phase peptide fractionation kit (Thermo Fisher Scientific) according to the manufacturer’s instructions. In brief, the reversed-phase fractionation spin column was washed twice with 300 μL of ACN, followed by conditioning twice with 300 μL of 0.1% TFA. The dried peptides were dissolved in 300 μL of 0.1% TFA, and then loaded onto the spin column. Additional washing was conducted with 300 μL H_2_O and 300 μL 5% ACN/0.1% TEA (triethylamine) to effectively remove unreacted TMT reagent. Following this, sequential elution was performed on the peptides with 300 μL of high-pH step-elution solutions. The solutions had increasing concentrations of ACN in 0.1% TEA, starting from 10% ACN, proceeding through 12.5%, 15%, 17.5%, 20%, 22.5%, 25%, and ending with 50% ACN. The resulting eight eluted fractions were dried using a Speed-Vac before LC-MS analysis.

### Liquid chromatography and mass spectrometry analysis

The fractionated peptides were re-suspended in 0.1% FA and separated using an in-house packed 20 cm × 75 μm internal diameter C18 column (1.9 μm ReproSil-Pur C18-AQ beads, Dr. Maisch GmbH, Germany) on a nanoflow Easy nLC 1200 UHPLC system (Thermo Fisher Scientific). The column was heated to 50°C using a home-made column heater. The flow rate was set at 300 nL/min. Buffer A and B were 0.1% FA in H_2_O and 0.1% FA in 80% acetonitrile, respectively. The separation gradient over a 120-min period was scheduled as: 2%-5% B in 1 min; 5%-32% B in 94 min; 32%-45% B in 15 min; 45%-65% B in 3 min; 65%-100% B in 1 min;100% B in 6 min. Samples were analyzed with a Q Exactive HF-X mass spectrometer (Thermo Fisher Scientific) equipped with a nanoflow ionization source. Data-dependent acquisition was performed in positive ion mode at a spray voltage of 2,300 V. The MS1 spectra was measured with a resolution of 120,000 @ m/z 200, an AGC target of 3e6, a maximum injection time of 50 ms and a mass range of 350 to 1,700 m/z. The data-dependent mode cycle was set to trigger MS2 scan on up to the top 20 most abundant precursors per cycle at an MS2 resolution of 45,000 @ m/z 200, an AGC target of 1e5, a maximum injection time of 120 ms, an isolation window of 1.0 m/z, an HCD (high collision dissociation) collision energy of 32, and a fixed first mass of 105.0 m/z. The dynamic exclusion time was set as 40 s and precursor ions with charge 1, 7, 8 and > 8 were excluded for MS2 analysis.

### Analysis of mass spectrometry data

MS raw files were searched against the UniProt database containing 20,588 human reference proteome sequences using MaxQuant^19^ (version 2.4.2.0). TMT 10-plex based MS2 reporter ion quantification was chosen with reporter mass tolerance set as 0.003 Da. The purities of TMT labeling channels were corrected according to the kit LOT number. Enzyme digestion specificity was set to Trypsin and maximum two missed cleavages were allowed. Carbamidomethyl cysteine was set as fixed modification. Oxidized methionine and protein N-term acetylation were set as variable modifications. The tolerances of first search and main search for peptides were set at 20 ppm and 4.5 ppm, respectively. A cut-off of 1% FDR was applied at the peptide and protein level.

Complete mass spectrometry data are shown in Supplementary Table 1 and the analyses were performed as previously described^10,18^. Proteins identified by two or more unique peptides were considered for the dataset. To normalize input levels for each sample, normalization was carried out by dividing all TMT ratios by the median of the ratios for false-positive (FP) proteins that should not be biotinylated by APEX2+SOPP3. The log_2_ value of each of these ratio values was then calculated. To determine the cutoff ratio for each comparison, a receiver operating characteristic (ROC) analysis was performed.

For comparison of APEX2+SOPP3 MAM labeling against omit light controls, APEX2+SOPP3 labeling against APEX2 controls, and APEX2+SOPP3 labeling against SOPP3 controls, true-positive (TP) proteins were literature-validated ER-mitochondria contact proteins (Supplementary Table 2) and false-positive (FP_A) proteins were known mitochondrial matrix proteins annotated by GO: 0005759 and not annotated for OMM (GO:0005741), IMS (GO:0005758) or IMM (GO:0005743). For each comparison, the proteins were ranked in descending order based on corresponding TMT log_2_ ratios in each replicate. To determine optimal cutoffs, the true positive rate (TPR) and false positive rate (FPR) were calculated at each possible TMT log_2_ ratio. TPR/FPR is defined as the fraction of TP/FP proteins above the TMT log_2_ ratio in that replicate. A ROC curve was then plotted for each comparison using these calculated TPR and FPR values. The optimal cutoff was set where TPR-FPR maximized. After applying these cutoffs to each comparison, resulting proteomic lists were intersected, yielding shared protein datasets of 944 and 1025 proteins for replicate 1 and replicate 2 respectively.

For comparison of APEX2+SOPP3 MAM labeling against cytosolic NES controls, a different set of false-positive (FP_B) proteins was used. These proteins were known cytosol-resident proteins as determined by GOCC (GO:0005829, but no annotations for membrane) in a previous study^10^. After filtering with the ROC cutoff as described above, the resulting lists from each replicate were overlapped, and a final list of 84 proteins was obtained and compiled in Supplementary Table 1.

For the Gene Ontology (GO) enrichment analysis, the final proteomic list for mam APEX2+SOPP3 was analyzed with the R package clusterProfiler^20^ (version 4.6.2), and the top 10 terms for GO cellular component (GOCC) are displayed. Additionally, the sub-cellular locations of these proteins were annotated using the GOCC and UniProt databases.

### Isolation of crude mitochondria and proteinase K protection assay

Mitochondria were extracted using Cell Mitochondria Isolation Kit (Beyotime Biotechnology, C3601) according to the manufacturer’s instruction. Freshly isolated crude mitochondria from HeLa cells were resuspended in mitochondria storage buffer (Beyotime Biotechnology, C3601). Samples were treated with proteinase K (Sigma-Aldrich, 0.1-10 μg/mL) for 30 min on ice to digest surface-exposed protein. The reaction was stopped by adding PMSF (2 mM) and sample buffer, followed with western blotting analysis.

### Western blotting analysis

Lysate from culture cells were harvested as previously descried^18^. Proteins were separated by SDS-PAGE and blotted onto polyvinylidene fluoride membranes. The following antibodies were used: mouse anti-V5 (Abclonal, AE017), rabbit anti-HA (Cell signaling technology, 3724S), rabbit anti-VDAC1 (Abcam, ab306581), rabbit anti-RMDN3 (Abclonal, A5820), rabbit anti-FTH1 (Abclonal, A1144), rabbit anti-TMX1 (Abclonal, A17219), rabbit anti-BCAP31 (Abclonal, A7056), rabbit anti-TOM20 (Sigma-Aldrich, HPA011562), rabbit anti-Tim23 (Proteintech, 11123-1-AP), mouse anti-HSP60 (Proteintech, 66041-1-Ig), mouse anti-α-Tubulin (Sigma-Aldrich, T9026), mouse anti-FLAG (Sigma-Aldrich, F1804), Streptavidin-HRP (Invitrogen, S911), HRP-Goat anti-Rabbit IgG (H+L) (Jackson ImmunoResearch, 111-035-045), HRP-Goat anti-Mouse IgG (H+L) (Jackson ImmunoResearch, 115-035-062).

### Immunostaining, confocal imaging, and data processing

Immunostaining in HeLa cells was performed as previously descried^18^. The following antibodies were used: mouse anti-V5 (Abclonal, AE017), rabbit anti-HA (Cell signaling technology, 3724S), Goat-Alexa Fluore 647-conjugated anti-mouse (Invitrogen, A32728), Goat-Alexa Fluore 488-conjugated anti-rabbit (Invitrogen, A11034) and Alexa Fluore 555-conjugated streptavidin (Invitrogen, S32355). The Annexin V-mCherry Apoptosis Detection Kit was a product of Beyotime Company (Shanghai, China, C1069S). The images were captured by Olympus spinning disk confocal microscopes with 60X objective.

Imaging data processing was performed in ImageJ (NIH) and cellSens (Olympus). 3D reconstruction of APEX2+SOPP3 mediated PL in HeLa cells MAM and cell surface (Supplementary Video 1, 2) were generated using Imaris x64 software (version 10.0, Bitplane). Stack images were first converted to an imaris file (.ims) using Imaris File Converter. 3D reconstruction of the confocal image was performed using the Surface rendering option in the Surpass view, with a thresholding method based on the Absolute Intensity of the signal. Movies were generated from Imaris x64 software (version 10.0, Bitplane).

### Flow cytometry analysis

For flow cytometric analysis of cell surface biotinylation, cells were washed with DPBS supplemented with quencher solution after illumination or H_2_O_2_ treatment. The cell pellets were resuspended in Stain Buffer (2% FBS in DPBS) with Rabbit anti-HA (Cell signaling technology, 3724S) antibody diluted at 1:500. All samples were incubated for 1 hour on ice protected from light. Cells were then extensively washed with DPBS and stained with Alexa Fluore 555-conjugated streptavidin (Invitrogen, S32355) and Goat-Alexa Fluore 488-conjugated anti-rabbit (Invitrogen, A11034) for 1 hour on ice prior analysis using CytoFLEX LX (BECKMAN COULTER) and FlowJo (10.4.0).

### Statistics

Data were analyzed with the Graphpad Prism 5 software using the unpaired two-tailed student’s-test. The plot error values were calculated by standard error of the mean (SEM). All data in this study were repeated for at least three times.

## Acknowledgements

We are grateful to the members of W.P. laboratory for technical supports. We also thank Dr. Dianqing (Dan) Wu from Yale University School of Medicine for the constructive suggestions in our manuscript preparation. This work was supported by the National Key Research and Development Plan of China (2018YFA0800203), the National Natural Science Foundation of China for Distinguished Young Scholars (31925014), the National Natural Science Foundation of China Key Program (32130033), the Key Research Program of Frontier Sciences, Chinese Academy of Sciences (ZDBS-LY-SM010), Shanghai Pilot Program for Basic Research-Chinese Academy of Sciences, Shanghai Branch (JCYJ-SHFY-2022-006) and Shanghai Science, Technology Innovation Action Plan for Basic Research Program (21JC1406300) and the CAS project for Young Scientists in Basic Research (YSBR-077), and the National Key Research and Development Plan of China (2022YFA1302902).

## Author contribution

W.P. conceived and supervised the project. D.Q., W.Z., Y.L. planed and performed most of the experiments. Q.L. performed mass spectrometry measurement under the supervision of H.Z. D.Q., W.Z., Y.L. analyzed data, made figures and models under the supervision of W.P. All authors contributed to data interpretation. W.P., D.Q. and W.Z. wrote the manuscript with input from all authors.

## Data and Code availability

The data associated with this study are in the article and the Supplementary information. The mass spectrometry proteomics data have been deposited to the ProteomeXchange Consortium via the PRIDE partner repository with the dataset identifier PXD052044. Additional data beyond that provided in the figures and Supplementary Information are available from the corresponding author on request. There is no original code generated in this study.

