## Supplementary information for "Photoactivated SOPP3 enables APEX2-mediated proximity labeling with high spatio-temporal resolution in live cells"

- **This PDF file includes:**

Supplementary Figure S1-6

Legends for Supplementary Table S1-3

Legends for Supplementary Video S1-2

- **Other Supplementary Information for this manuscript include the following:**

Supplementary Table S1-3 as .xls file

Supplementary Video S1-2 as .mp4 file

**Supplementary Figures**

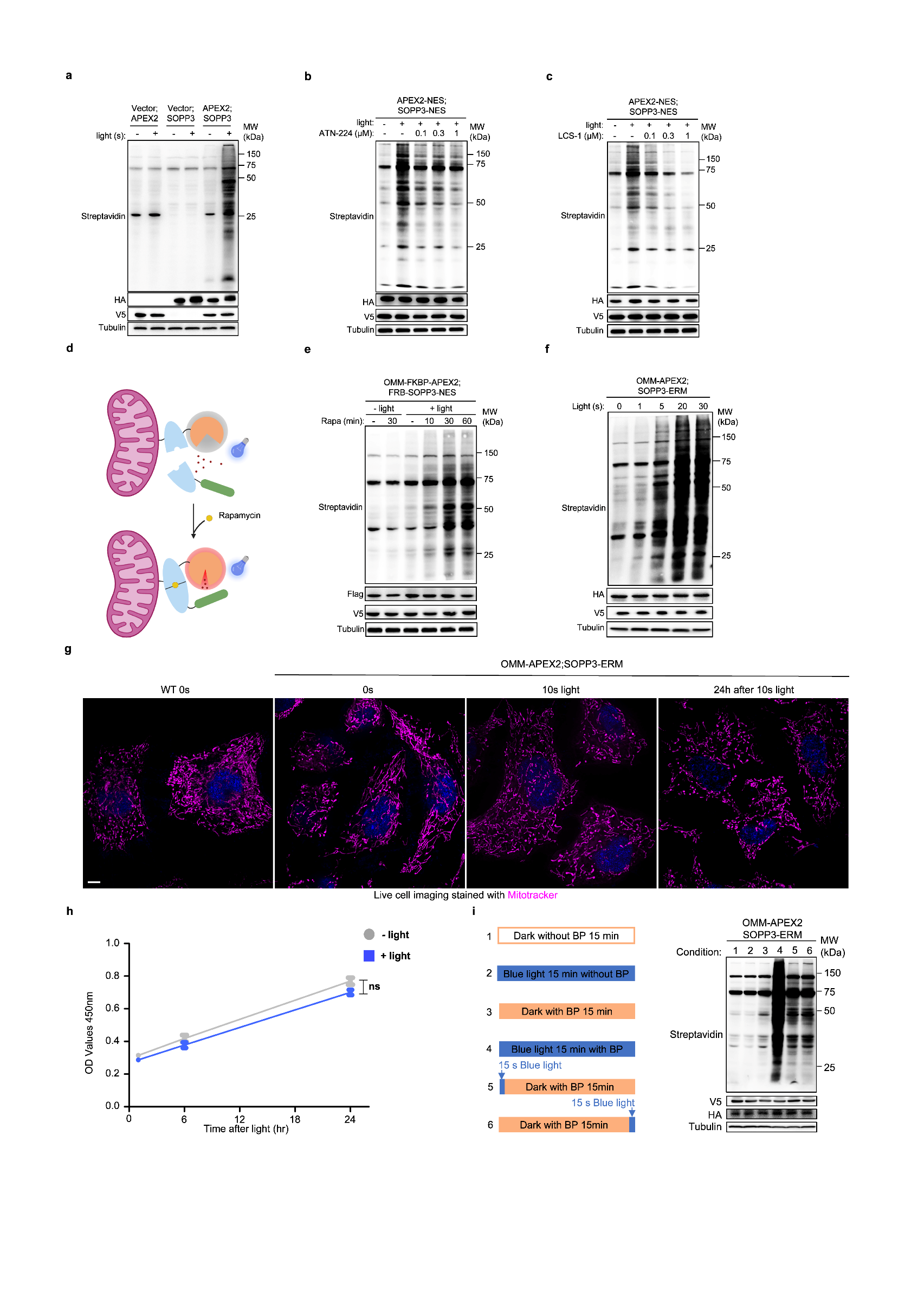
**Figure. S1**

**Fig. S1. Additional data related to characterization on APEX2+SOPP3-mediated proximity labeling**

**a** Evaluation on the efficiency of photo-proximity labeling mediated by APEX2+SOPP3 compared with APEX2 or SOPP3 alone. Streptavidin blot showed labeling efficiency, anti-V5 and anti-HA showed expression level of APEX2 and SOPP3. Anti-α-tubulin was used as loading control in this study.

**b,c** Evaluation on the efficiency of photo-proximity labeling mediated by APEX2+SOPP3 upon SOD1 inhibitor. HeLa cells, after transfection of constructs indicated in figure, were treated with ATN-224 (**b**) or LCS-1 (**c**) in indicated concentration for 16 hours. Cells were then treated with 500 μM BP for 1 hour and subsequently illuminated with blue light. Streptavidin blot showed labeling efficiency, anti-V5 and anti-HA showed expression level of APEX2-NES and SOPP3-NES.

**d-e** Schematic (d) and evaluation (e) on proximity dependency of APEX2+SOPP3 mediated PL via rapamycin-induced FKBP-FRB interaction. Anti-V5 and anti-Flag indicated expression level of OMM-FKBP-APEX2 and FRB-SOPP3-NES respectively. Illumination time, 10 seconds.

**f** APEX2-SOPP3-mediated proximity labeling in time course. HeLa cells transiently transfected with APEX2 and SOPP3, targeted to mitochondria and ER constructs respectively, were treated with 500 μM BP for 1 hour and then illuminated for indicated times. Biotinylation was detected by western blotting analysis. Streptavidin blot showed labeling efficiency, anti-V5 and anti-HA showed expression level of OMM-APEX2 and SOPP3-ERM.

**g** Confocal live cell imaging of mitochondria morphology after APEX2+SOPP3-mediated proximity labeling. HeLa cells expressing APEX2 and SOPP3, targeted to OMM and ERM respectively, were illuminated for indicated times, and then stained with Mitotracker to probe morphology of mitochondria in live cells, followed with live cell confocal imaging. Scale bar, 5 μm.

**h** Cell viability analysis on HeLa cells expressing APEX2+SOPP3 targeted to Mito/ER and treated with or without 10 seconds illumination. N.S., not significant (*P*=0.8187). Data were shown as mean ± SEM (n=3). Statistical significance was determined using unpaired two-tailed Student’s *t*-test with no adjustments.

**i** APEX2+SOPP3-mediated proximal labeling terminated when the blue light illumination stopped. The cells expressing APEX2/SOPP3 targeted to OMM/ERM were activated as conditions indicated in the figure, and then harvested for streptavidin blotting analysis. Streptavidin blot showed labeling efficiency, anti-V5 and anti-HA showed expression level of OMM-APEX2 and SOPP3-ERM.

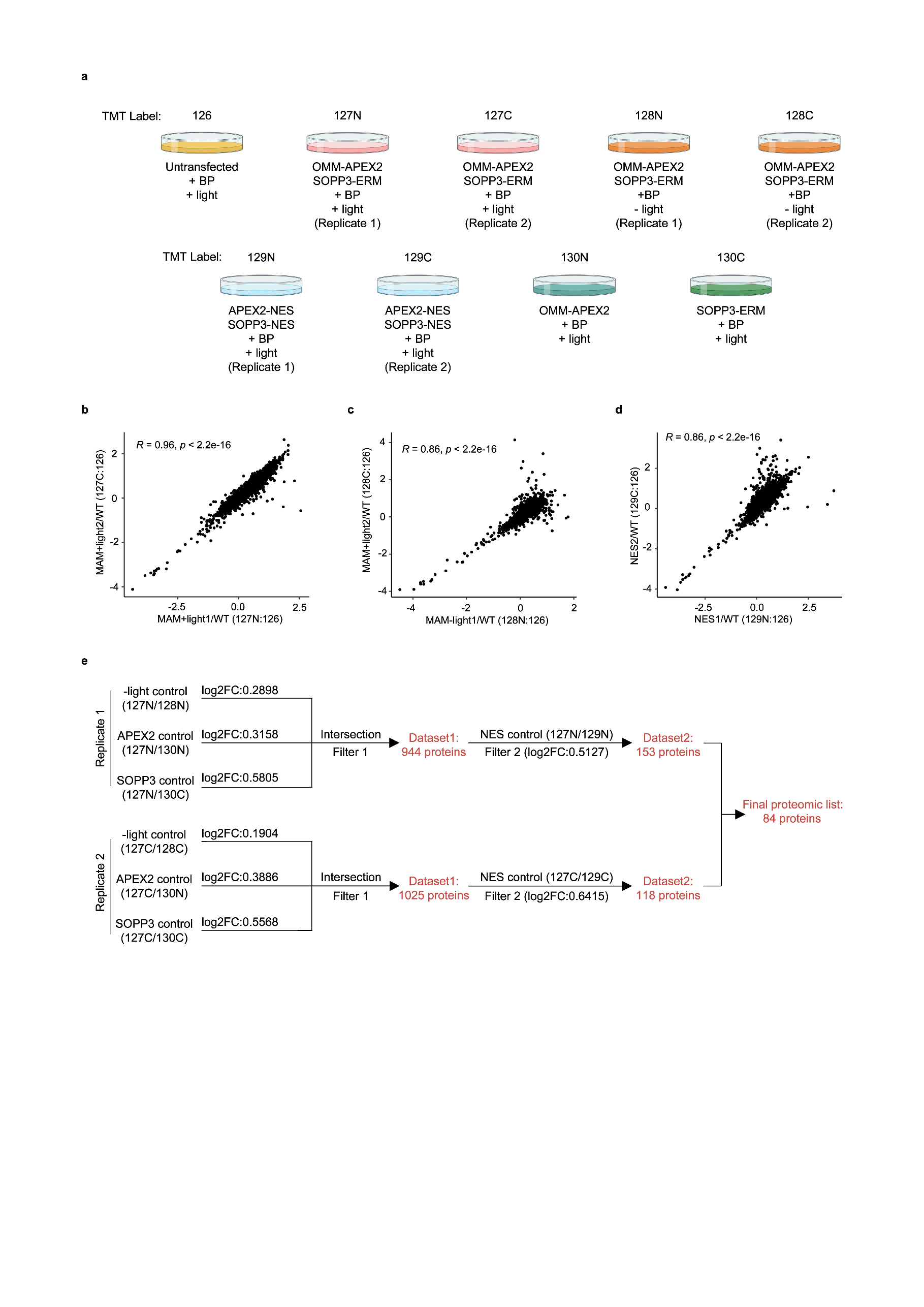
**Figure. S2**

**Fig. S2. Additional analysis on proteomic data**

**a** Experimental design and labeling conditions for tandem mass tag (TMT)-based proteomics.

**b-d** Scatterplots of log2 ratios for replicates of APEX2+SOPP3-mediated proximal labeling on MAM APEX2+SOPP3 + light, MAM APEX2+SOPP3 - light, NES APEX2+SOPP3 + light. MAM (APEX2 and SOPP3, targeted to OMM and ERM respectively), NES (APEX2 and SOPP3, both expressed in cytosol).

**e** Filtering scheme for mass spectrometric data analysis.

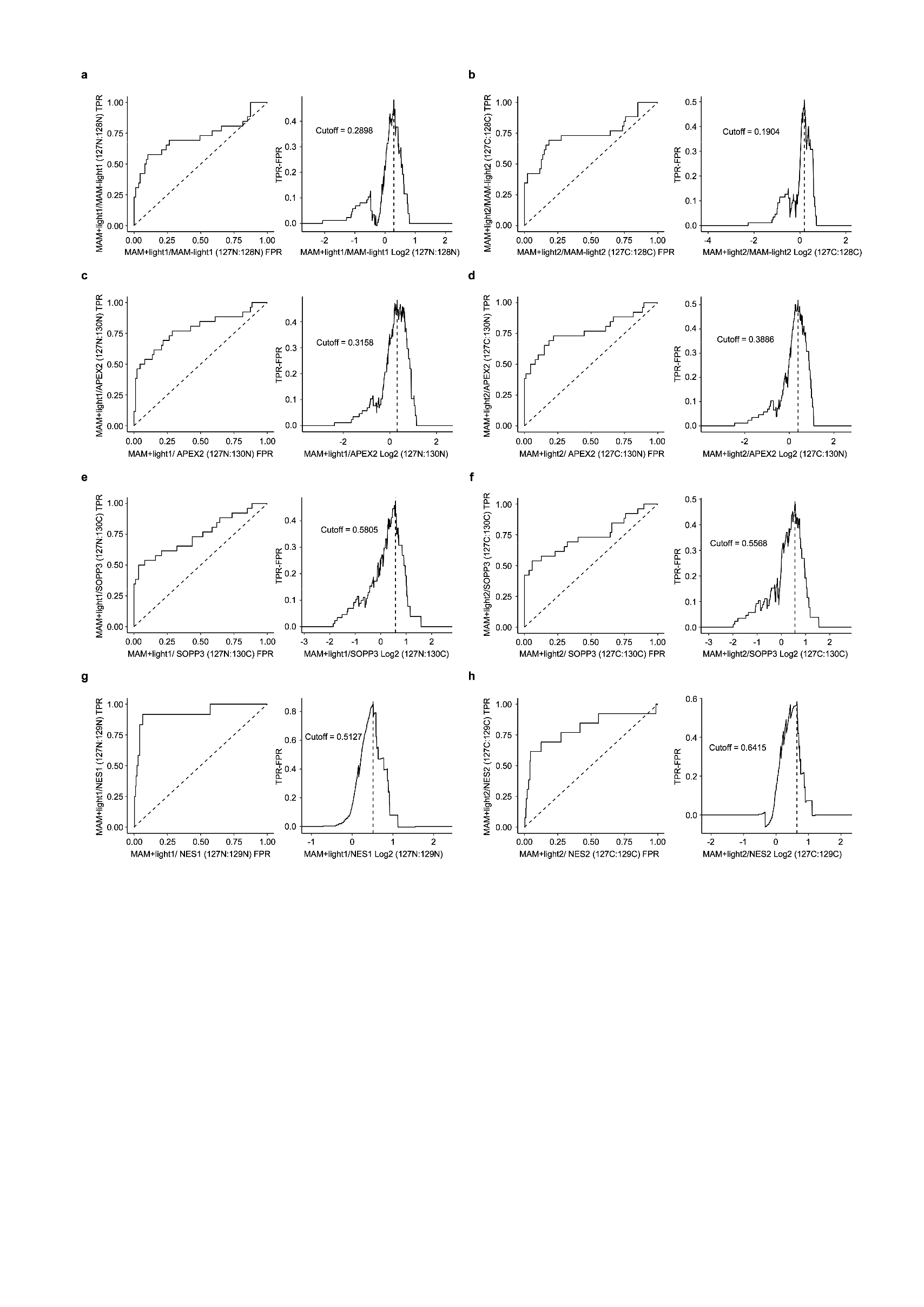
**Figure. S3**

**Fig. S3. Additional analysis on proteomic data**

**a-b** Receiver operator curve (ROC) analysis of MAM proteome identified by APEX2+SOPP3-mediated proximity labeling method. ROC curves of MAM-1 +/- light (a) and MAM-2 +/- light (b) were used to determine cut-off ratio. The optimal cut-off was set where TPR-FPR maximized.

**c-d** Receiver operator curve (ROC) analysis of MAM proteome identified by APEX2+SOPP3-mediated proximity labeling method. ROC curves of MAM-1 + light / APEX2 + light (c) and MAM-2 + light / APEX2 + light (d) were used to determine cut-off ratio. The optimal cut-off was set where TPR-FPR maximized.

**e-f** Receiver operator curve (ROC) analysis of MAM proteome identified by APEX2+SOPP3-mediated proximity labeling method. ROC curves of MAM-1 + light / SOPP3 + light (e) and MAM-2 + light / SOPP3 + light (f) were used to determine cut-off ratio. The optimal cut-off was set where TPR-FPR maximized.

**g-h** Receiver operator curve (ROC) analysis of MAM proteome identified by APEX2+SOPP3-mediated proximity labeling method. ROC curves of MAM-1 + light / NES-1 + light (g) and MAM-2 + light / NES-2 + light (h) were used to determine cut-off ratio. The optimal cut-off was set where TPR-FPR maximized.

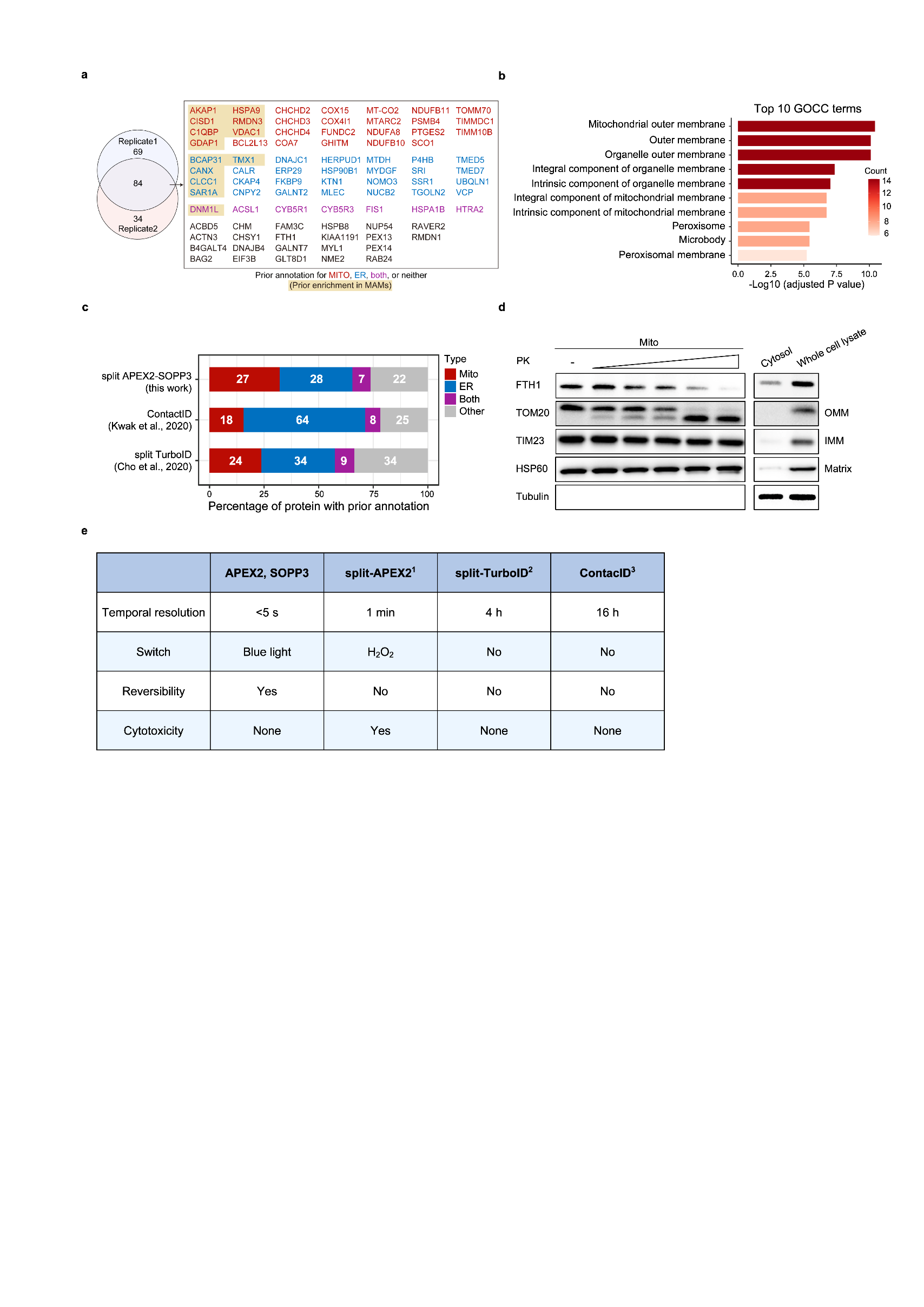
**Figure. S4**

**Fig. S4. Additional analysis and validation of proteomic data**

**a** Venn diagram of MAM proteome obtained from **fig 1e**. Proteins with annotated subcellular localization on mitochondria (Mito), ER, on both Mito and ER, or neither, are labeled in red, blue, purple or black. Proteins, enriched in MAMs known from literature, are highlighted in yellow.

**b** Top 10 Gene Ontology Cellular Component (GOCC) terms for APEX2+SOPP3-mediated proximity labeling targeted to MAM proteome (84 proteins).

**c** Specificity analysis on proteomic datasets generated using APEX2+SOPP3-mediated proximity labeling compared to previously published datasets. Bar graph shows the percentage of each proteome with identified proteins classified as 1) prior mitochondria annotation only, 2) ERM annotation only, 3) both mitochondria and ERM annotation or 4) other annotation. Each bar is labeled with the size of the proteome.

**d** Biochemical validation of FTH1 distribution on MAM. Crude mitochondria (Mito) isolated from HeLa cells were subjected to Proteinase K (PK) digestion. FTH1 was analyzed by western blotting. Outer mitochondrial membrane (OMM) protein TOM20, inner mitochondrial membrane (IMM) protein TIM23, mitochondrial matrix protein HSP60 and cytosolic protein α-Tubulin were used as control.

**e** Comparison of APEX2+SOPP3 with split proximity labeling methods in literature^1-3^.

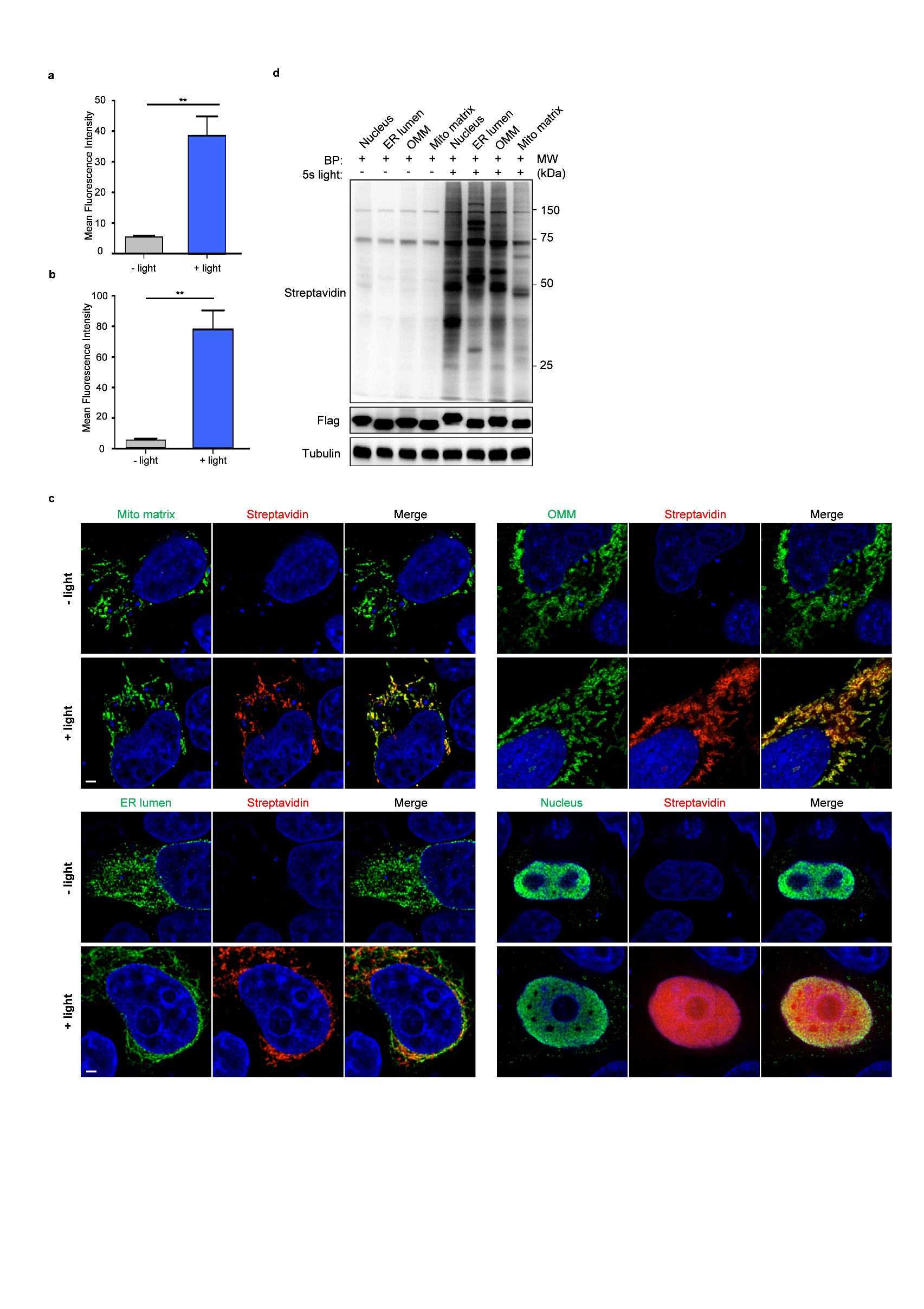
**Figure. S5**

**Fig. S5. Additional data related to chimeric APEX2-SOPP3-mediated proximity labeling**

**a,b** Evaluation on efficiency of ERM-targeted (a) or cell surface-targeted (b) chimeric APEX2-SOPP3-mediated proximity labeling via Alexa Fluore 555-conjugated streptavidin signal quantification. For **a** (***P*=0.0066) and **b** (***P*=0.0047), data were shown as mean±SEM (n=3). Statistical significance was determined using unpaired two-tailed Student’s t-test with no adjustments.

**c-d** Confocal fluorescence imaging of photo-PL via chimeric APEX2-SOPP3 targeted to various subcellular compartments (**c**). Chimeric APEX2-SOPP3-ERM and biotinylation signals was visualized by anti-HA antibody and Alexa Fluor 555-conjugated streptavidin respectively. Evaluation on photo-PL efficiency in **c** via western blot (**d**). Anti-Flag blot indicated expression level of chimeric APEX-SOPP3 targeted to various subcellular compartments. Illumination time, 5 seconds. Scale bar, 10 μm.

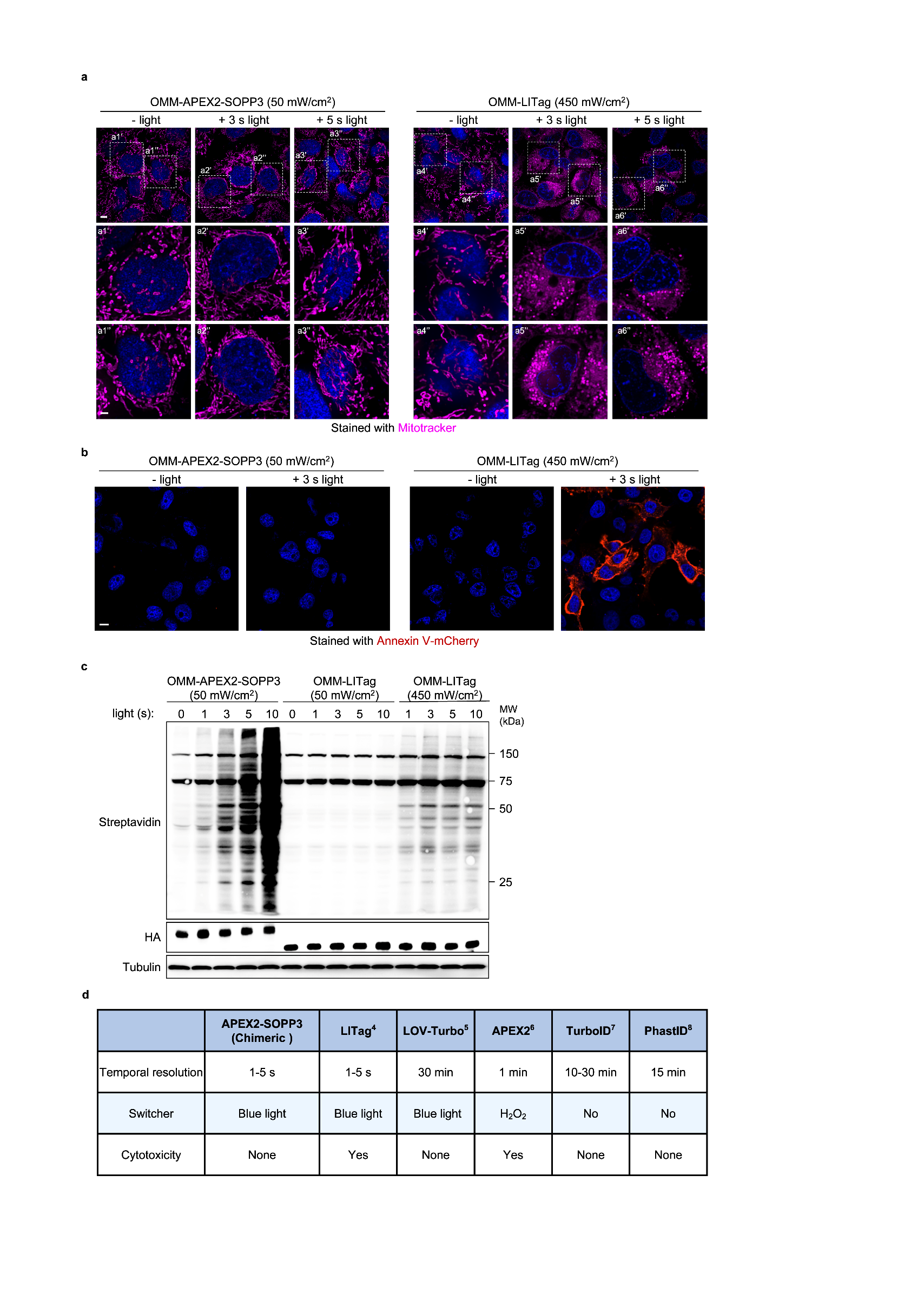
**Figure. S6**

**Fig. S6. Comparison of chimeric APEX2-SOPP3 with LITag-mediated proximity labeling**

**a** Confocal live cell imaging analysis on the perturbation of mitochondria morphology via chimeric APEX2-SOPP3 or LITag-mediated proximity labeling. HeLa cells transfected with chimeric APEX2-SOPP3 or LITag, targeted to outer membrane of mitochondria, were treated with 500 μM BP for 1 h and then illuminated for 0 ,3 or 5 seconds via different light power indicated in the figure. Cells were stained with Mitotracker to probe morphology of mitochondria via confocal live cell imaging. Scale bar, 5 μm. Zoomed images from boxed region. Scale bar, 2 μm.

**b** Cytotoxicity evaluation on APEX2-SOPP3 or LITag-mediated proximity labeling. HeLa cells transfected with chimeric APEX2-SOPP3 or LITag, targeted to outer membrane of mitochondria, were treated with 500 μM BP for 1 h and then were illuminated for 3 seconds via different light power indicated in the figure, and then stained with Annexin V to detect apoptosis level. Scale bar, 10 μm.

**c** Comparison on proximity labeling efficiency of chimeric APEX2-SOPP3 and LITag. HeLa cells transfected with chimeric APEX2-SOPP3 or LITag, targeted to outer membrane of mitochondria, were treated with 500 μM BP for 1 h and then illuminated for indicated times. Biotinylation was analyzed by western blotting. Streptavidin blot showed labeling efficiency, anti-HA showed expression level of chimeric OMM-APEX2-SOPP3 and OMM-LITag. Anti-α-tubulin was used as a loading control.

**d** Comparison of chimeric APEX2-SOPP3 with current PL technologies^4-8^.

**Legends for Supplementary Tables**

**Supplementary information, Table S1 (Separated file).** Data collection of mass spectrometry proteomics

**Supplementary information, Table S2 (Separated file).** List of true positive (TP) and false positive (FP) for proteomics data analysis

**Supplementary information, Table S3 (Separated file).** List of plasmids and oligos used in this study

**Legends for Supplementary Videos**

**Supplementary information, Video S1 (Separated file). 3D reconstruction of APEX2+SOPP3-mediated proximity labeling on ER-Mito contact sites in HeLa cell.**

The ‘Surface’ tool in Imaris software was used to create a 3D rendering of each channel of confocal images (links to Fig. 1f). Contact area algorithm was further performed to determine the interface between OMM-APEX2 (cyan) and ERM-SOPP3 (green). The interfaces (yellow) largely overlapped with biotinylation signals (red), indicating the accuracy of APEX2-SOPP3-mediated proximity labeling.

**Supplementary information, Video S2 (Separated file). 3D reconstruction of APEX2+SOPP3-mediated proximity labeling on cell-cell interface.**

The ‘Surface’ tool in Imaris software was used to create a 3D rendering of each channel of confocal images (links to Fig. 1j). Contact area algorithm was further performed to determine the interface between HeLa cell expressing APEX2-TM (cyan) and neighboring HeLa cell expressing SOPP3-TM surface (green). The interfaces were rendered in yellow, which was largely overlapped with biotinylation signals rendered in red.
